# Conformations of the Human Immunodeficiency Virus (HIV-1) Envelope Glycoproteins in Detergents and Styrene-Maleic Acid Lipid Particles (SMALPs)

**DOI:** 10.1101/2023.03.01.530731

**Authors:** Rong Zhou, Shijian Zhang, Hanh T. Nguyen, Haitao Ding, Althea Gaffney, John C. Kappes, Amos B. Smith, Joseph G. Sodroski

**Affiliations:** Department of Cancer Immunology and Virology, Dana-Farber Cancer Institute, Boston, MA 02215, USA; Department of Microbiology, Harvard Medical School, Boston, MA 02115, USA; Department of Medicine, University of Alabama at Birmingham, AL 35294, USA; Birmingham Veterans Affairs Medical Center, Research Service, Birmingham, AL 35294, USA; Department of Chemistry, University of Pennsylvania, Philadelphia, PA 19104, USA; Current affiliation: Department of Cardiology, The Second Affiliated Hospital of Shanxi Medical University, Taiyuan, Shanxi, China

**Keywords:** Virus, membrane, Env, solubilization, Cymal-5, DIBMA, SMA, default conformation, structure, detergent

## Abstract

The mature human immunodeficiency virus (HIV-1) envelope glycoprotein (Env) trimer, which consists of non-covalently associated gp120 exterior and gp41 transmembrane subunits, mediates virus entry into cells. The pretriggered (State-1) Env conformation is the major target for broadly neutralizing antibodies (bNAbs), whereas receptor-induced downstream Env conformations elicit immunodominant, poorly neutralizing antibody (pNAb) responses. To examine the contribution of membrane anchorage to the maintenance of the metastable pretriggered Env conformation, we compared wild-type and State-1-stabilized Envs solubilized in detergents or in styrene-maleic acid (SMA) copolymers. SMA directly incorporates membrane lipids and resident membrane proteins into lipid nanodiscs (SMALPs). The integrity of the Env trimer in SMALPs was maintained at both 4°C and room temperature. By contrast, Envs solubilized in Cymal-5, a non-ionic detergent, were unstable at room temperature, although their stability was improved at 4°C and after incubation with the entry inhibitor BMS-806. Envs solubilized in ionic detergents were relatively unstable at either temperature. Comparison of Envs solubilized in Cymal-5 and SMA at 4°C revealed subtle differences in bNAb binding to the gp41 membrane-proximal external region (MPER), consistent with these distinct modes of Env solubilization. Otherwise, the antigenicity of the Cymal-5- and SMA-solubilized Envs was remarkably similar, both in the absence and presence of BMS-806. However, both solubilized Envs were recognized differently from the mature membrane Env by specific bNAbs and pNAbs. Thus, detergent-based and detergent-free solubilization at 4°C alters the pretriggered membrane Env conformation in consistent ways, indicating that loss of Env association with the membrane results in default state(s).

**IMPORTANCE:** The human immunodeficiency virus (HIV-1) envelope glycoproteins (Envs) in the viral membrane mediate virus entry into the host cell and are targeted by neutralizing antibodies elicited by natural infection or vaccines. Detailed studies of membrane proteins rely on purification procedures that allow the proteins to maintain their natural conformation. In this study, we show that a styrene-maleic acid (SMA) copolymer can extract HIV-1 Env from a membrane without the use of detergents. The Env in SMA is more stable at room temperature than Env in detergents. The purified Env in SMA maintains many but not all of the characteristics expected of the natural membrane Env. Our results underscore the importance of the membrane environment to the native conformation of HIV-1 Env. Purification methods that bypass the need for detergents could be useful tools for future studies of HIV-1 Env structure and its interaction with receptors and antibodies.

## INTRODUCTION

Human immunodeficiency virus (HIV-1) entry is mediated by the envelope glycoprotein (Env) trimer, a Class I viral fusion protein composed of three gp120 exterior subunits and three gp41 transmembrane subunits (1–3). In HIV-1 infected cells, Env is synthesized in the endoplasmic reticulum as an ∼856-amino acid precursor that trimerizes and undergoes signal peptide cleavage and the addition of high-mannose glycans (4–7). In the Golgi compartment, the gp160 Env precursor is cleaved by host furin-like proteases into the gp120 and gp41 subunits and is further modified by the addition of complex carbohydrates (8–12). These mature Envs are selectively incorporated into virions (8).

Single-molecule fluorescence resonance energy transfer (smFRET) experiments indicate that, on virus particles, the Env trimer exists in three conformational states (States 1 to 3) (13). The metastable pretriggered Env conformation (State 1) predominates on virions from primary HIV-1 strains; upon interaction with the receptors, CD4 and CCR5 or CXCR4, Env undergoes transitions to lower-energy states (States 2 and 3) (13–15). Receptor-induced transitions in gp41 result in the interaction of the N-terminal fusion peptide with the target cell membrane and in the formation of a six-helix bundle that drives the fusion of viral and cell membranes (16–24).

In natural HIV-1 infection, Env strain variability, heavy glycosylation and conformational flexibility contribute to HIV-1 persistence by diminishing the binding and elicitation of neutralizing antibodies (25–29). The gp160 Env precursor, which samples multiple conformations, and disassembled Envs (shed gp120, gp41 six-helix bundles) elicit high titers of poorly neutralizing antibodies (pNAbs) that fail to recognize the mature pretriggered (State-1) Env trimer (30–35). After years of infection, a minority of HIV-1-infected individuals generate broadly neutralizing antibodies (bNAbs), most of which recognize the cleaved pretriggered (State-1) Env conformation (13, 36–43). Six categories of bNAbs that target different Env epitopes have been identified (Table 1) (27,29,44,45). Although passively administered monoclonal bNAbs are protective in animal models of HIV-1 infection (46–49), suggesting their potential utility in prophylaxis, bNAbs have not been efficiently and consistently elicited by current vaccine candidates (50–54).

**Table 1.** Antibodies recognizing HIV-1 Env epitopes.

**Broadly neutralizing antibodies (bNAbs)**
| <u>Antibody</u> | <u>Epitope</u> |
| --- | --- |
| 2G12 | gp120 outer domain glycans |
| PGT121 | gp120 V3 glycans |
| PG9<br>PG16<br>PGT145 | gp120 V1/V2 quaternary (trimer apex) |
| VRC01<br>VRC03 | gp120 CD4-binding site (CD4BS) |
| PGT151<br>35O22 | gp120-gp41 interface |
| 4E10<br>10E8 | gp41 membrane-proximal external region (MPER) |

**Poorly neutralizing antibodies (pNAbs)**
| <u>Antibody</u> | <u>Epitope</u> |
| --- | --- |
| 19b | gp120 V3 |
| F105 | gp120 CD4-binding site (CD4BS) |
| 902090 | gp120 V2 linear |
| 17b | gp120 CD4-induced (CD4i) |
| F240 | gp41 disulfide loop (Cluster I) |

Differences in the antigenicity, glycosylation and conformation of soluble Env trimers used as immunogens and the pretriggered (State-1) membrane Env have been observed (12,55–59). Given the requirement for bNAbs to recognize conserved, conformation-specific and often glycan-dependent elements on the pretriggered (State-1) Env (13,36–43), even small differences from the native State-1 Env might affect immunogen efficacy. Both the conformation and the glycan composition of the State-1 Env depend upon association with the membrane (12,57,60–65). These considerations recommend the need to study HIV-1 Envs in native membrane environments.

In preparation for their biochemical, biophysical and structural characterization, membrane proteins are often solubilized in detergents to allow purification. In some cases, the purified proteins can be reconstituted into a phospholipid membrane environment such as proteoliposomes or nanodiscs (66, 67). For metastable membrane proteins such as HIV-1 Env, detergent solubilization could lead to irreversible effects on conformation. Indeed, Env conformations resembling State 2, which has been suggested to represent a default conformation (68, 69), have been observed for detergent-solubilized HIV-1 Envs, including those reconstituted into proteoliposomes or nanodiscs (58,70–73). To allow membrane protein purification and characterization while bypassing completely the need for detergent solubilization, styrene-maleic acid (SMA) copolymers have been developed that directly solubilize lipid membranes and resident membrane proteins into 9-15-nm discoidal nanoparticles (SMA lipid particles or SMALPs) (74–80). In some cases, membrane proteins in SMALPs have been shown to be thermostable compared to the detergent-solubilized proteins (80, 81). The cryo-electron microscopy (cryo-EM) structure of the multidrug exporter AcrB in SMALPs revealed a remarkably well-ordered lipid bilayer associated with the transmembrane domains of the protein (82). Here, we compare the stability and antigenicity of HIV-1 Envs solubilized using SMA versus detergents (Cymal-5 or SDS/deoxycholate). Such studies provide insights into the important contribution of protein-membrane interactions to the maintenance of a native Env conformation.

## RESULTS

### Antigenicity of a State-1-stabilized HIV-1_AD8_ Env variant on the surface of expressing cells

A previous study described HIV-1_AD8_ Env mutants with increased State-1 stability, improved precursor cleavage, and a higher content of lysine residues potentially useful for crosslinking (83). One of these HIV-1_AD8_ variants, 2-4 RM6 E, will be used throughout this study and, in some cases, will be compared with the wild-type (wt) HIV-1_AD8_ Env (see Material and Methods for a more detailed description of these Envs). To provide a standard for the antigenicity of a native, State-1-enriched membrane Env, we evaluated the recognition of cleaved and uncleaved 2-4 RM6 E Env on the surface of expressing A549 cells by panels of pNAbs and bNAbs (Fig. 1, Table 1). As expected (43, 71), the pNAbs preferentially recognized the 2-4 RM6 E gp160 Env but not the cleaved gp120 and gp41 Envs. By contrast, the bNAbs precipitated the cleaved gp120 and gp41 Envs, indicating their ability to bind the mature Env trimer on the cell surface. The antigenicity profile of the cell-surface 2-4 RM6 E Env is similar to that previously reported for the wt HIV-1_AD8_ Env (43).

**FIG 1.**
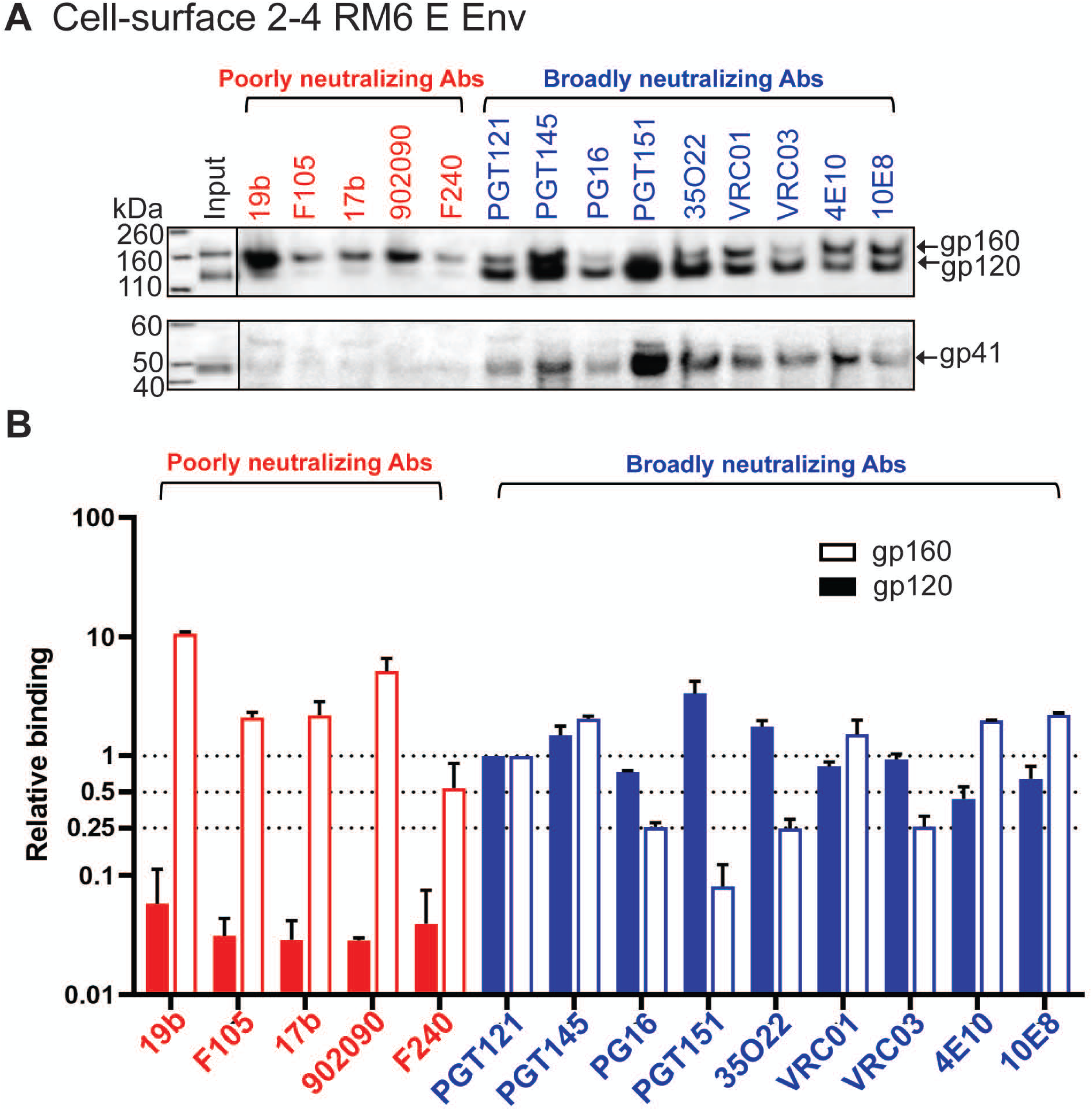
Antibody recognition of the 2-4 RM6 E Env on the cell surface. (A) A549 cells were induced with doxycycline to express 2-4 RM6 E Env. The cells were detached from the tissue-culture plates with 5 mM EDTA in 1x PBS. After pelleting and resuspension, the cells were aliquoted and incubated with the indicated antibodies. After washing, the cells were lysed in NP-40 lysis buffer and the cell lysates were incubated with Protein A-Sepharose beads. The precipitated antibody-Env complexes were Western blotted with a goat anti-gp120 antiserum (upper panel) or the human 4E10 anti-gp41 antibody (lower panel). The experiment was repeated twice and a typical result is shown. (B) The gp160 and gp120 bands in A were quantified using Fiji Image J (NIH). The intensities of the gp160 and gp120 bands precipitated by each antibody were normalized to those of the respective gp160 and gp120 bands precipitated by the PGT121 antibody. The means and standard deviations of the results of two independent experiments are shown.

### Antigenicity of wt and 2-4 RM6 E Envs in a cell membrane preparation

Preparation of cell membranes is often used as an initial step in the purification of membrane proteins, as cell nuclei, mitochondria and cytosolic proteins are removed. To evaluate the potential effect of membrane preparation on Env conformation, we examined the antigenicity of the wt HIV-1_AD8_ and the 2-4 RM6 E Envs in membranes purified from expressing A549 cells (Fig. 2). We evaluated membrane Env antigenicity in the absence and presence of BMS-806, a small-molecule HIV-1 entry inhibitor that decreases Env transitions from State 1 and thereby stabilizes the pretriggered conformation (13,43,59,84–86). Briefly, after incubation with a panel of pNAbs and bNAbs, the purified cell membranes were lysed in an NP-40 buffer and the Env-antibody complexes were captured on Protein A-Sepharose beads and analyzed by Western blotting. The 2-4 RM6 E Env was proteolytically processed more efficiently than the wt HIV-1_AD8_ Env, as expected (83), resulting in a relative increase in the cleaved 2-4 RM6 E Env in the Input samples (Fig. 2A). Most pNAbs efficiently recognized the wt and 2-4 RM6 E gp160 Envs. The addition of BMS-806 slightly reduced gp160 binding by the F105 pNAb against the CD4 binding site and the 17b pNAb against a CD4-induced (CD4i) gp120 epitope; both epitopes are near the BMS-806 binding site (87). Recognition of the cleaved Envs by the pNAbs was generally inefficient; however, the 19b anti-V3 pNAb and to a lesser extent the 17b CD4i pNAb recognized the wt and 2-4 RM6 E gp120 glycoproteins. This binding of the 19b and 17b pNAbs to gp120 was inhibited by BMS-806; moreover, when normalized to the Input gp120, the recognition of the 2-4 RM6 E gp120 by the 19b and 17b pNAbs was reduced compared with that of the wt HIV-1_AD8_ Env (Fig. 2B, left panels). These observations suggest that the exposure of V3 and CD4i epitopes may result from the spontaneous sampling of more open State 2/3-like conformations by the Envs on the purified cell membranes. Spontaneous exposure of gp120 V3 and CD4i epitopes on intact HIV-1_AD8_ virions has recently been observed (88) and is consistent with the conformational flexibility of virion Envs documented by smFRET studies (13,59,85). Two representative bNAbs, PGT121 against a V3-glycan epitope and PGT145 against the trimer apex, recognized the cleaved wt and 2-4 RM6 E Envs efficiently in the absence or presence of BMS-806. Thus, the antibody binding profiles of the wt HIV-1_AD8_ and 2-4 RM6 E Envs in the purified membranes correspond to those observed for these Envs on the cell and viral surfaces (Fig.1 and references 43 and 88).

**FIG 2.**
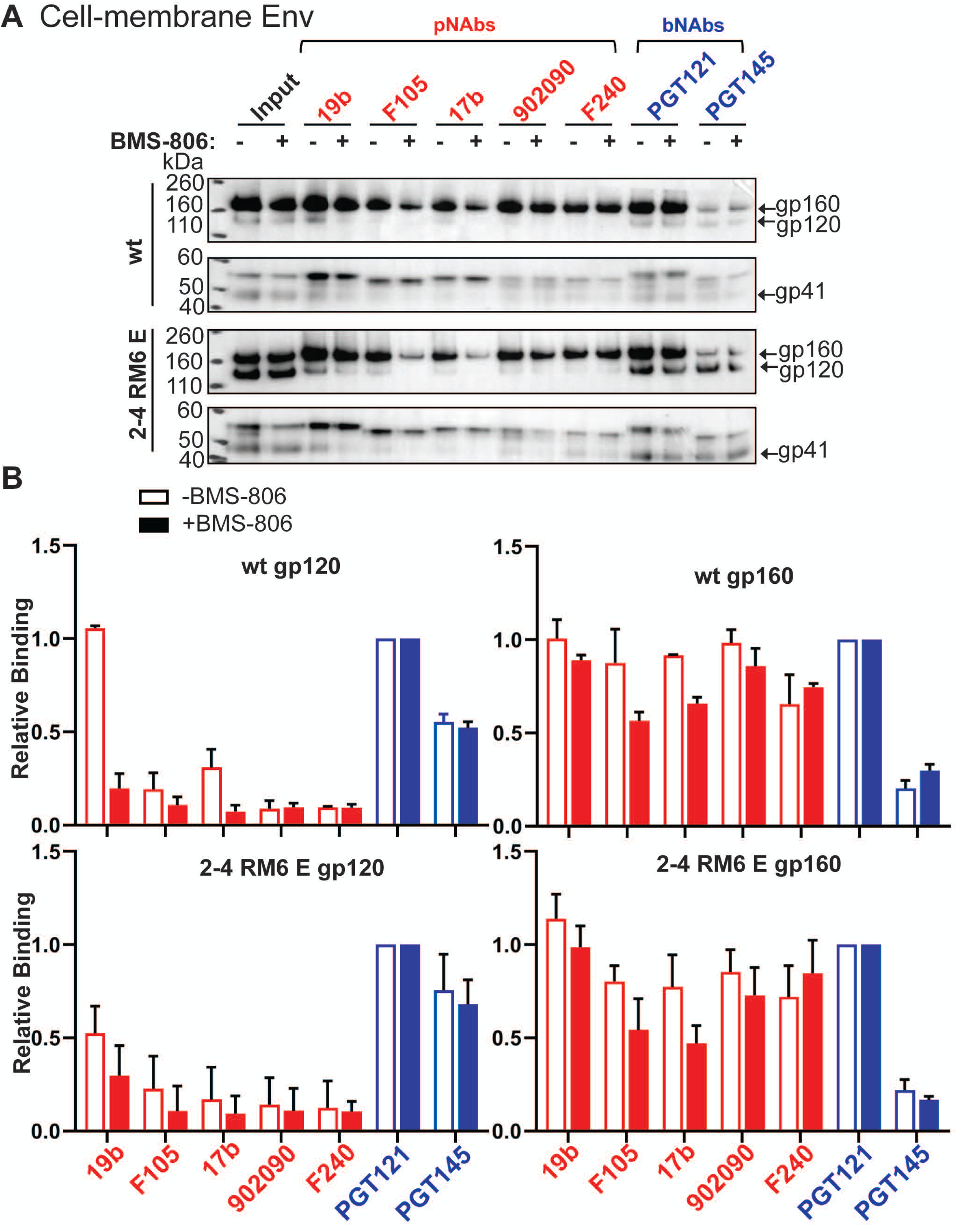
Antigenicity profiles of HIV-1 Envs on purified cell membranes. (A) A549 cells were induced with doxycycline to express the wt HIV-1_AD8_ Env or the 2-4 RM6 E Env. Cell membranes were purified as described in Materials and Methods and incubated with the indicated antibodies for 1 h at 4°C, with or without 10 µM BMS-806. The membranes were lysed with 1% NP-40 with or without BMS-806 for 5 minutes on ice, and then incubated with Protein A-Sepharose beads for 1 h at 4°C. The precipitated proteins were subjected to Western blotting with a goat anti-gp120 antiserum or the human 4E10 anti-gp41 antibody. (B) The intensities of the gp120 and gp160 bands in A were measured in Fiji Image J (NIH). The gp120 and gp160 band intensity for each antibody was normalized to those of the respective gp120 and gp160 bands of the PGT121 samples. The means and standard deviations from two independent experiments are shown.

### Comparison of the ability of different amphipathic copolymers to extract HIV-1 Env from membranes

Although SMALPs have proven to be of great utility for membrane protein solubilization and characterization (74–82), they do have limitations. SMALPs need to be maintained at low concentrations of divalent cations and at pH values above 7.5 to remain in solution (77, 80). Likely due to the aromatic styrene groups in the SMA copolymer, SMALPs absorb ultraviolet light, interfering with protein concentration measurements at 280 nm (77). Efforts have been made to overcome these limitations by replacing maleic acid with other hydrophilic moieties (e.g., glycerol, glucosamine) or supplanting styrene with diisobutyrene (Fig. 3) (89, 90).

**FIG 3.**
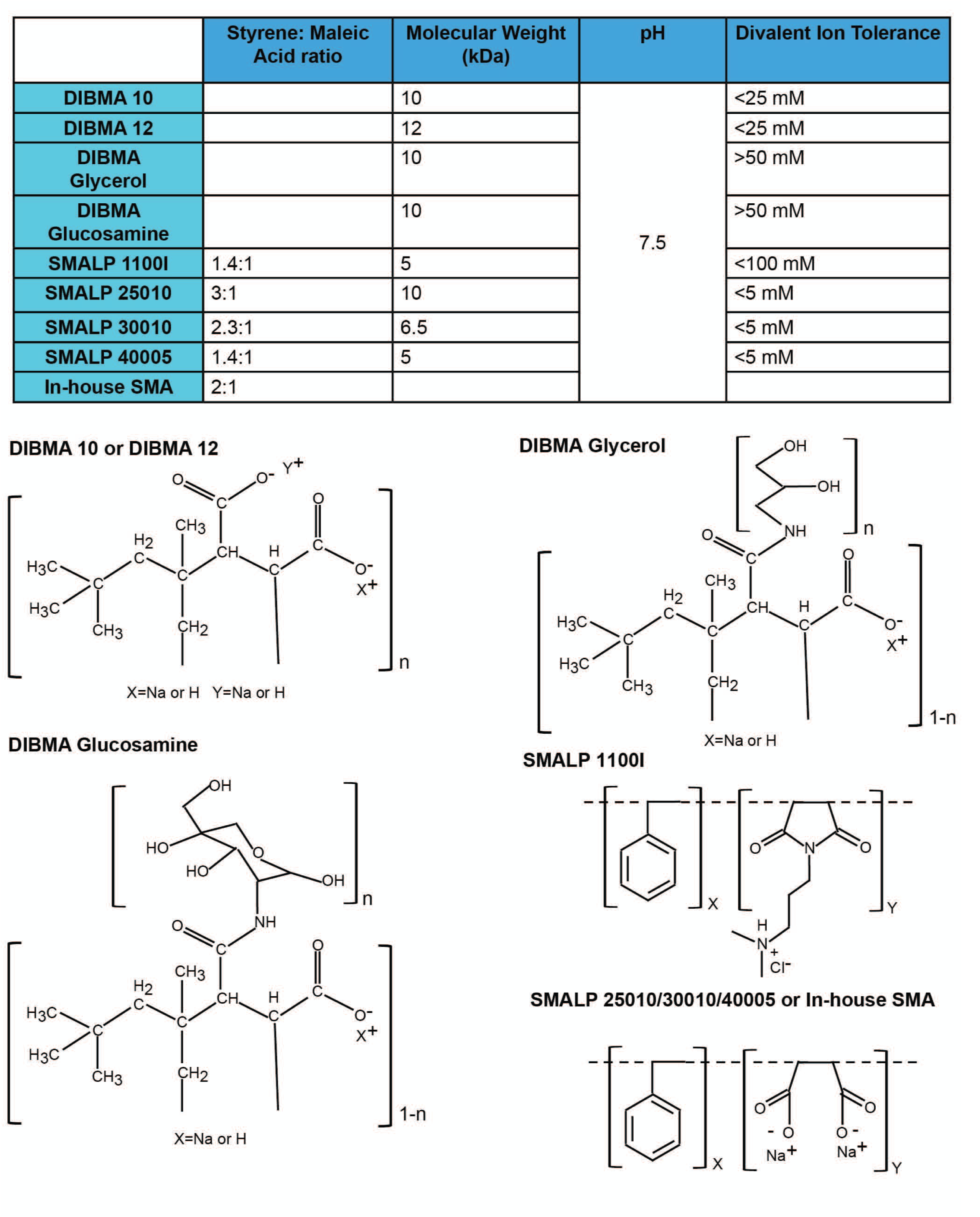
Properties of amphipathic copolymers for membrane protein solubilization. The structures and properties of the SMA and DIBMA variants used in this study are shown.

We compared commercially available preparations of SMA and diisobutyrene-maleic acid (DIBMA) variants with an in-house preparation of SMA (see Materials and Methods). The efficiency with which these amphipathic copolymers extracted a crosslinked, State-1-stabilized Env, AE.2, from cell membranes was compared. The AE.2 Env is closely related to the 2-4 RM6 E Env, but has one additional State-1-stabilizing change and one additional lysine substitution for potential crosslinking (see Materials and Methods) (83). Cell membranes prepared from A549 cells expressing the His_6_-tagged AE.2 Env were crosslinked with DTSSP. The crosslinked membrane was divided into equal parts, which were used for the extraction of Env by the different amphipathic copolymers. The solubilized Envs were purified on Ni-NTA beads and subjected to SDS-PAGE and silver staining (Fig. 4A). The highest yield of Env was obtained with our in-house SMA, with slightly lower Env yields obtained with DIBMA 10, DIBMA 12 and SMA 11001. The SMA 11001, which unlike the other SMA variants has a dimethylaminopropylamine (DMAPA) group, extracted a higher proportion of uncleaved gp160 and crosslinked gp120-gp41 Env than the other SMA copolymers. These results indicate that the ratio and the nature of the hydrophobic and polar groups in the copolymer can influence the yield and maturity of the extracted Envs.

**FIG 4.**
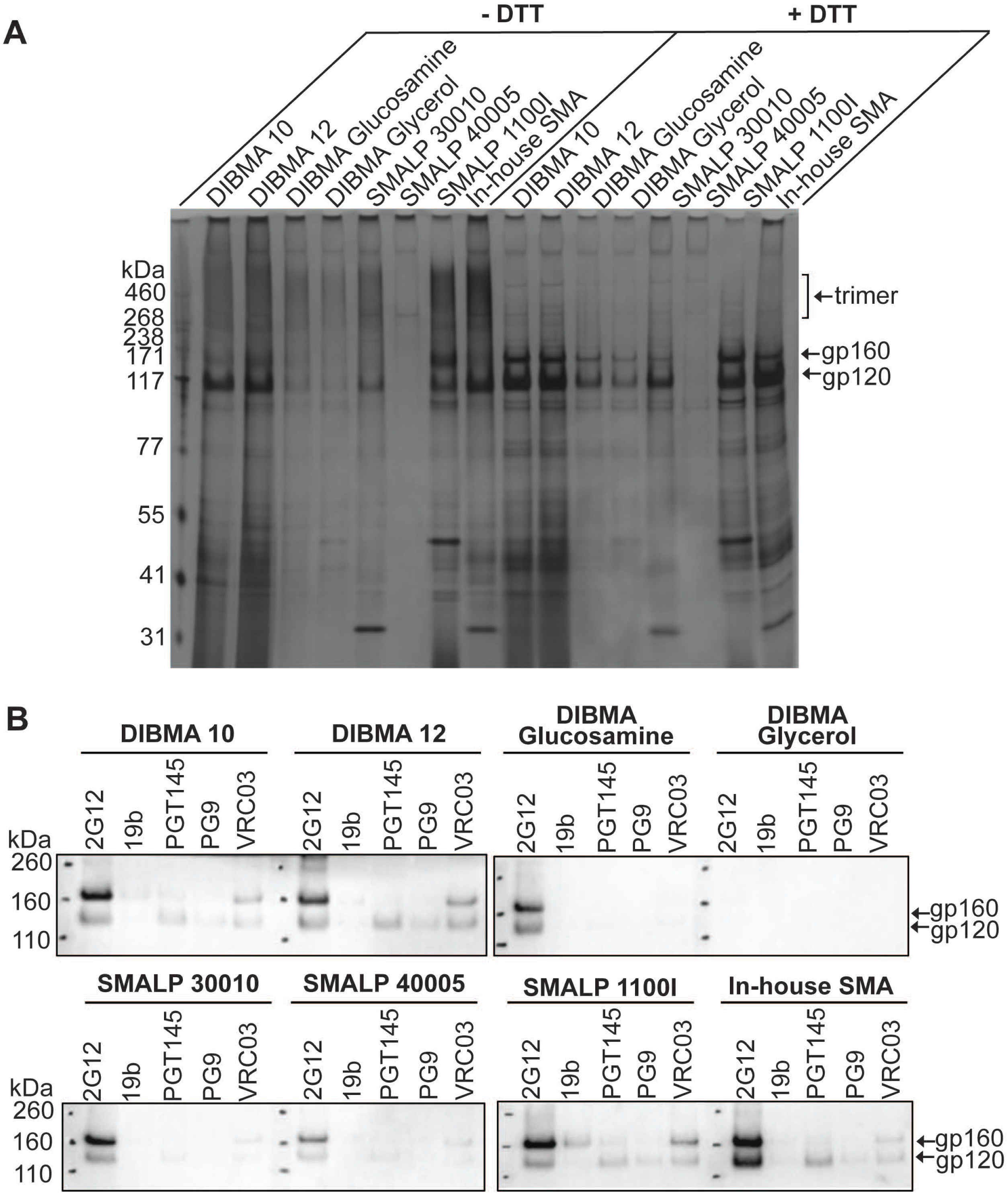
Yield and antigenicity of the AE.2 Env solubilized in different amphipathic copolymers. A549 cells were induced with doxycycline to express the AE.2 Env. Cell membranes were purified, crosslinked with 0.35 mM DTSSP and extracted with the indicated amphipathic copolymers. (A) Eluates from the Ni-NTA beads were analyzed by SDS-PAGE under non-reducing (-DTT) or reducing (+DTT) conditions, followed by silver staining. (B) A small panel of antibodies was used to assess the antigenicity of the AE.2 Env eluted from the Ni-NTA beads. The 2G12 bNAb recognizes gp120 outer domain glycans in a manner that is relatively independent of Env conformation. The 19b pNAb recognizes a gp120 V3 epitope. The PGT145, PG9 and VRC03 bNAbs exhibit some preference for the pretriggered (State-1) Env conformation. The AE.2 Env precipitated by the antibodies was captured on Protein A-Sepharose beads and subjected to Western blotting with a goat anti-gp120 antiserum. The results of a typical experiment are shown.

We next examined the antigenic profiles of the crosslinked and purified AE.2 Envs extracted with the different amphipathic copolymers. The ability of four bNAbs (2G12, PGT145, PG9 and VRC03) and the 19b pNAb to recognize the solubilized AE.2 Envs was evaluated. PGT145, PG9 and VRC03 exhibit some preference for State-1 Envs; 2G12 recognizes gp120 outer domain glycans in a manner that is less dependent on the Env conformational state (13,43,68,69). Also, PGT145 and PG9 preferentially recognize cleaved Envs (13,68,69,91). Differences in the yield of Env influenced the amounts of Env precipitated by the antibodies (Fig. 4B). For the instances where bNAb recognition was detectable, the overall pattern of bNAb recognition was similar for all copolymers. The 2G12 bNAb recognized most of the solubilized Envs in proportion to the yields. On the other hand, recognition of the purified AE.2 Envs by the State-1 preferring bNAbs was relatively weak, indicating that the pretriggered Env conformation was only inefficiently preserved in the amphipathic copolymer complexes. With the exception of the uncleaved AE.2 Env extracted by SMALP 11001, the purified AE.2 Envs were inefficiently recognized by the 19b pNAb. As the yield of antigenically similar, cleaved Env trimers extracted by our in-house SMA was at least as good as that of any of the other copolymers, we used our in-house SMA for the following studies.

### Stability of solubilized Env trimers at different temperatures

Some membrane proteins in SMALPs or DIBMA nanodiscs have been suggested to be more stable than when solubilized in detergents (79,81,82,92–94). We compared the stability of the wt HIV-1_AD8_ and 2-4 RM6 E Env trimers solubilized in Cymal-5 (a nonionic detergent), SMA and RIPA buffer (which contains two ionic detergents, SDS and deoxycholate). Both Envs were extracted from expressing cells slightly more efficiently by Cymal-5 and RIPA than by SMA (see Input in Fig. 5A). To evaluate the stability of the solubilized Envs, we took advantage of the His_6_ tag at the gp41 C-terminus, which allows the Envs to be precipitated using Ni-NTA beads. The ratio of the gp120:gp160 Envs in the precipitated samples provides an indication of the stability of the Env trimer in solution (43). In this manner, Env trimer stability was measured at either 4°C or room temperature for 1 and 16 h (Fig. 5A and B). Under all conditions tested, the wt and 2-4 RM6 E Envs were less stable in RIPA buffer than in Cymal-5 or SMA. The addition of BMS-806 during the solubilization only minimally improved the stability of the Envs in RIPA buffer; the positive effect of BMS-806 on RIPA-solubilized Envs was observed only during the 1-h experiment at 4°C. Apparently, the instability of the RIPA-solubilized Env trimers at room temperature or for longer incubation periods precluded the possibility of detecting a stabilizing effect of BMS-806.

**FIG 5.**
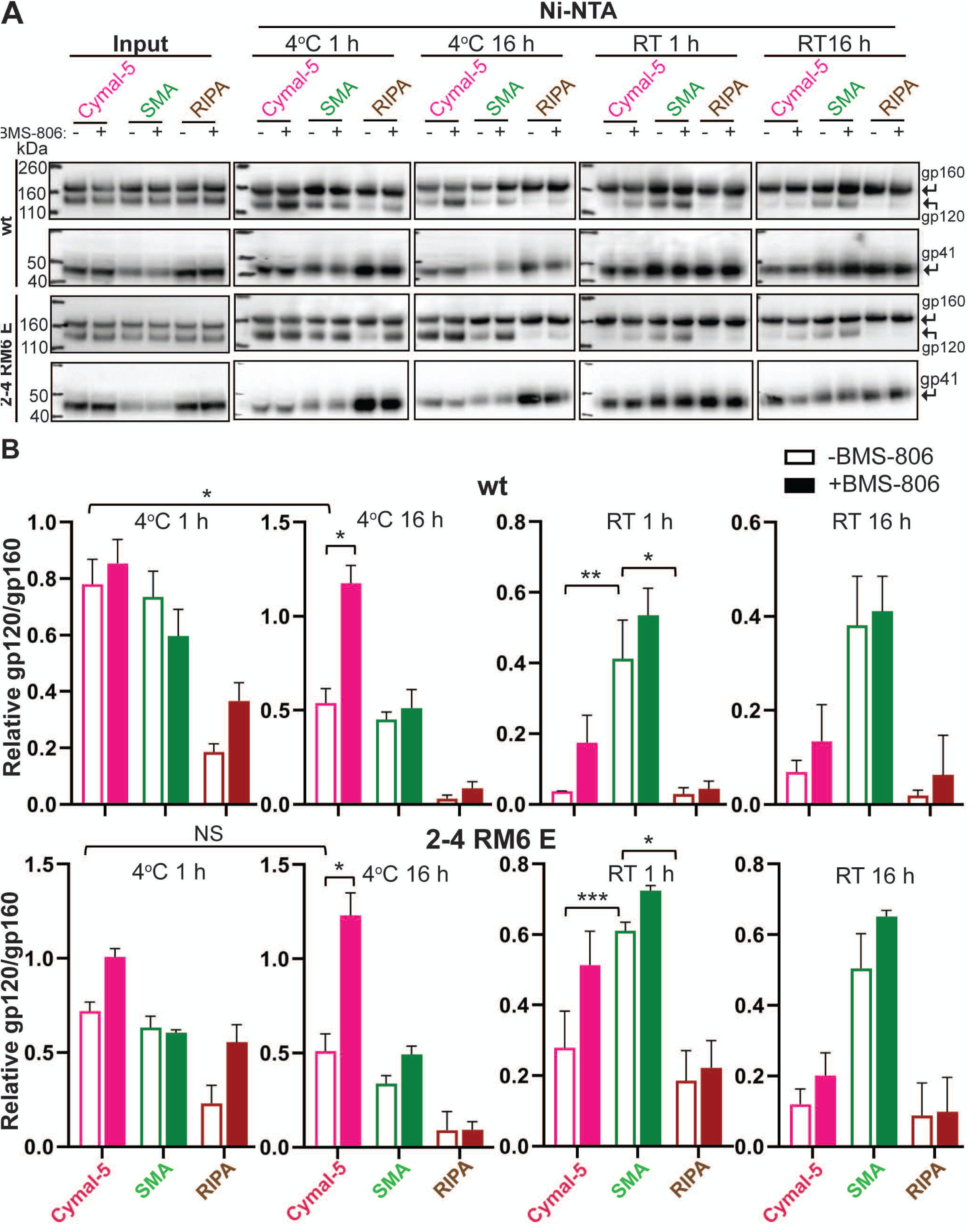
Stability of Env solubilized in Cymal-5, SMA or RIPA at different temperatures. To evaluate the integrity of the cleaved Env trimer under different solubilization conditions, we compared the relative efficiencies of gp120 and gp160 precipitation by Ni-NTA beads, which bind the His_6_ tags at the carboxyl termini of the Envs. (A) A549 cells were induced with doxycycline to express the wt HIV-1_AD8_ and 2-4 RM6 E Envs, both of which have His_6_ tags at the carboxyl terminus. Forty-eight hours after induction, the cells were lysed in 1% Cymal-5, 1% SMA or RIPA buffer in the absence or presence of 10 µM BMS-806 for 5 minutes on ice. After pelleting cell debris, a sample of the supernatant was saved as “Input.” The remainder of the cleared supernatants was incubated at the indicated times and temperatures with Ni-NTA beads by end-over-end rotation. For Cymal-5, SMA and RIPA samples, the beads were washed with 60 bed volumes of the corresponding wash buffers (20 mM Tris-HCl (pH 8.0), 100 mM (NH_4_)_2_SO_4_, 1 M NaCl, 30 mM imidazole, 1% Cymal-5); (20 mM Tris-HCl (pH 8.0), 100 mM (NH_4_)_2_SO_4_, 1 M NaCl, 30 mM imidazole, 0.005% Cymal-6); or RIPA buffer. After washing, the samples captured on the Ni-NTA beads were Western blotted with a goat anti-gp120 antiserum (upper panels) or the human 4E10 anti-gp41 antibody (lower panels). RT – room temperature. (B) Quantification of the gp120 and gp160 band intensity in A was performed in Fiji Image J (NIH). The stability of the Env trimer was calculated by the formula (gp120/gp160)_Ni-NTA_ + (gp120/gp160)_Input_. The means and standard deviations derived from two independent experiments are shown. The data were compared with a Student’s t test. Two-tailed P values are shown, *: P<0.05; **: P<0.01; ***: P<0.001.

The wt and 2-4 RM6 E Envs solubilized in SMA or Cymal-5 exhibited comparable stability during a 1-h incubation at 4°C in the absence of BMS-806; the stability of both Envs solubilized in Cymal-5 but not in SMA was enhanced by BMS-806 during a long (16-h) incubation at 4°C. Notably, at room temperature in the absence of BMS-806, both the wt and 2-4 RM6 E Envs in SMALPs were more stable than these Envs in Cymal-5 or RIPA buffer. The addition of BMS-806 did not significantly increase the stability of Envs in SMALPs at either temperature. Blue Native gel analysis revealed that at room temperature, the 2-4 RM6 E Env in SMALPs largely comprises higher-order oligomers consistent with trimers, whereas a substantial fraction of the 2-4 RM6 E Env in Cymal-5 consists of lower-molecular-weight forms (Fig. 6). Thus, compared with detergent-solubilized Envs, Envs in SMALPs maintained better trimer integrity during room temperature incubation.

**FIG 6.**
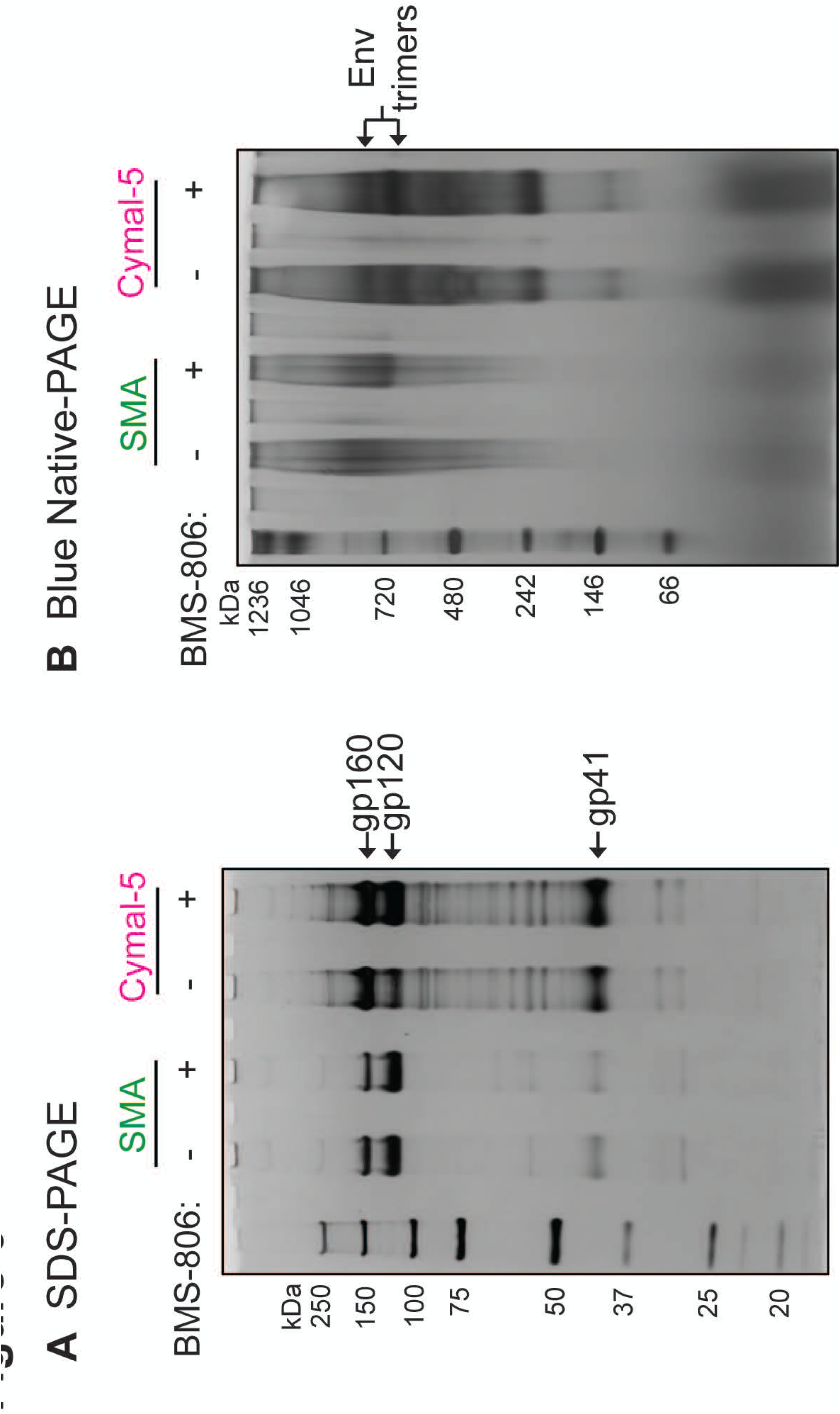
Oligomeric composition of the 2-4 RM6 E Env in Cymal-5 and SMA at room temperature. Membranes were prepared from A549 cells induced with doxycycline to express the 2-4 RM6 E Env. The 2-4 RM6 E Env was extracted at room temperature with 1% SMA or 1% Cymal-5. The extracted Envs were purified on Ni-NTA beads at room temperature and analyzed by SDS-PAGE (A) or Blue Native-PAGE (B), followed by silver staining. Note that BMS-806 treatment shifts the Env trimer population to a form that migrates faster on Blue Native gels.

### Antibody binding profile of the 2-4 RM6 E Env in Cymal-5 or SMALPs

The above observations indicate that solubilization in SMA and detergents can exert different effects on the stability of the Env trimers, raising the possibility that solubilization could influence Env antigenicity. To examine this possibility, lysates of A549 cells expressing the 2-4 RM6 E Env were prepared using 1% Cymal-5 or 1% SMA. Initially, we attempted to precipitate the solubilized Env from these cell lysates, but found that the presence of uncomplexed SMA in the lysates non-specifically interfered with immunoprecipitation by all antibodies tested (data not shown). This problem was remedied by partially purifying the 2-4 RM6 E Env in the cell lysates using Ni-NTA beads. After eluting the 2-4 RM6 E Env from the Ni-NTA beads, the antigenicity of the solubilized Env was evaluated by immunoprecipitation by a panel of pNAbs and bNAbs (Fig. 7A and B). In parallel experiments, BMS-806 was added to the cells prior to lysis and maintained in the cell lysates during the initial stage of immunoprecipitation.

**FIG 7.**
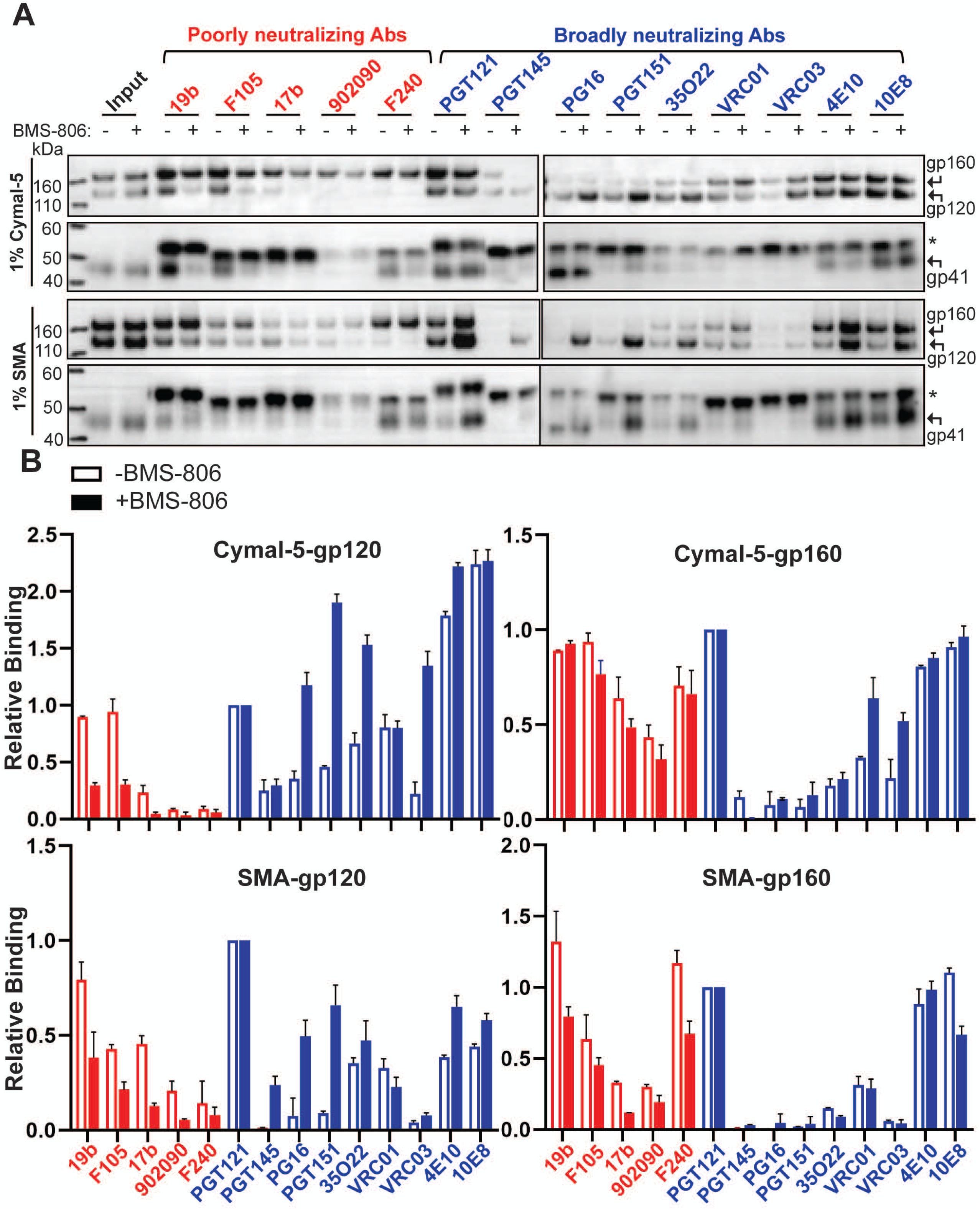
Antigenicity of the 2-4 RM6 E Env after Ni-NTA purification from cell lysates. (A) A549 cells were induced with doxycycline to express the 2-4 RM6 E Env. Forty-eight hours later, the cells were lysed with either 1% Cymal-5 or 1% SMA in the absence or presence of 10 µM BMS-806. The 2-4 RM6 E Env was partially purified by Ni-NTA affinity chromatography, aliquoted and incubated with the indicated antibodies together with Protein A-Sepharose beads for 1 h at 4°C. An aliquot without added antibody served as the Input sample. The Input sample and precipitated proteins were Western blotted with a goat anti-gp20 antiserum (upper panels) and the human 4E10 anti-gp41 antibody (lower panels). The asterisk marks the position of the heavy chains of the antibodies used for precipitation. (B) Quantification of the intensity of the gp120 and gp160 bands in A was performed in Fiji Image J (NIH). The intensities of the gp120 and gp160 bands were normalized to those of the respective gp120 and gp160 bands precipitated by the PGT121 antibody. The means and standard deviations derived from two independent experiments are shown.

In the absence of BMS-806, the cleaved 2-4 RM6 E Env in Cymal-5 was precipitated by all the bNAbs, although relatively weakly by the PGT145, PG16 and VRC03 bNAbs. Precipitation of the cleaved 2-4 RM6 E Env in SMALPs by bNAbs was generally weaker than that seen in Cymal-5, but showed a qualitatively similar pattern. Both Cymal-5- and SMA-solubilized cleaved 2-4 RM6 E Envs were recognized by the 19b and F105 pNAbs and, for the SMA-solubilized Envs, by the 17b pNAb as well. Both solubilized Envs contained populations that apparently shed gp120 and were precipitated by the F240 pNAb, which recognizes a Cluster I epitope on the gp41 ectodomain (95). These observations indicate that the conformations of both Cymal-5- and SMA-solubilized cleaved 2-4 RM6 E Envs differ from that of the cleaved cell-surface or membrane 2-4 RM6 E Env. In particular, the conformational differences between the solubilized cleaved Envs and the membrane cleaved Env are manifest in weaker recognition of the former Envs by the State-1-preferring PGT145, PG9 and VRC03 bNAbs and stronger recognition by V3, CD4BS and CD4i pNAbs. Thus, without BMS-806, the 2-4 RM6 E Env in Cymal-5 micelles or SMALPs readily samples conformations other than State 1.

It is noteworthy that in the absence of BMS-806, the antigenic profiles of the cleaved 2-4 RM6 E Envs solubilized in Cymal-5- and SMA are well correlated (Fig. 8A). The most significant differences between these solubilized Envs relate to recognition by the 4E10 and 10E8 bNAbs against the membrane-proximal external region (MPER) of gp41 (Fig. 8A); such differences may result from local changes in MPER structure associated with the distinct modes of Env solubilization employed by Cymal-5 and SMA. Recognition of other epitopes on the Cymal-5- and SMA-solubilized Envs was strongly correlated (Fig. 8B). Thus, in the absence of BMS-806 or other ligands, different means of solubilizing the 2-4 RM6 E Env result in similar conformational states.

**FIG 8.**
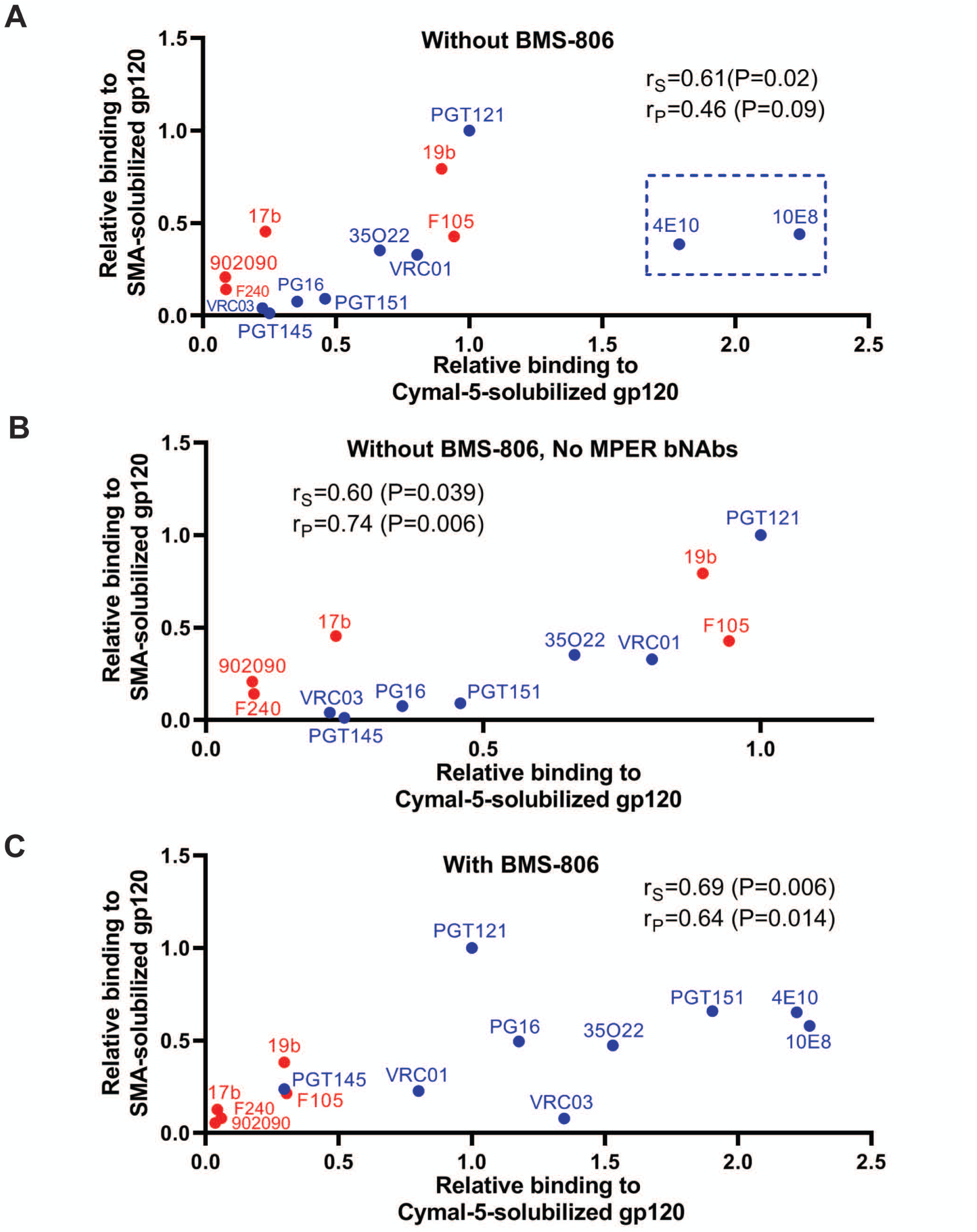
Comparison of the 2-4 RM6 E Env conformation in Cymal-5 and SMA. The correlation is shown between the antigenicities of the Cymal-5-solubilized and SMA-solubilized cleaved 2-4 RM6 E Envs in the absence (A,B) or presence (C) of 10 µM BMS-806. The pNAbs are designated in red and the bNAbs in blue. The Spearman rank correlation coefficient (r_S_), the Pearson r (r_P_) and two-tailed P values are shown. In A, note the outlier values for the two bNAbs (4E10 and 10E8) directed against the gp41 MPER (highlighted by the box). In B, the correlation is shown for the antibodies other than the MPER-directed 4E10 and 10E8 anti-gp41 antibodies.

The addition of BMS-806 changed the antigenic profile of the cleaved 2-4 RM6 E Env in Cymal-5 and SMALPs (Fig. 7A and B). Recognition of the cleaved 2-4 RM6 E Env by the PG16, PGT151 and 35O22 bNAbs (and by the PGT145 bNAb in the case of SMA) was increased in the presence of BMS-806. Recognition of the cleaved Env in Cymal-5 and SMALPs by the 19b, F105, 17b and F240 pNAbs was decreased by BMS-806. Thus, both forms of solubilized Env demonstrated an antigenic profile closer to that of the cell-surface Env when BMS-806 was present. Nonetheless, the relatively weak binding of the PGT145 and VRC03 bNAbs and the residual binding of the 19b and F105 pNAbs to the cleaved 2-4 RM6 E Env in Cymal-5 and SMALPs indicate conformational differences from the mature membrane Env. Similar to our observation in the absence of BMS-806 (Fig. 8A), we observed that the antigenic profiles of the Cymal-5- and SMA-solubilized 2-4 RM6 E Envs correlated in the presence of BMS-806 (Fig. 8C). Apparently, distinct modalities that extract Env from its membrane environment generate BMS-806-bound Envs in similar, non-State-1 conformations.

As expected (33,43,71), the uncleaved gp160 forms of both Cymal-5 and SMA-solubilized 2-4 RM6 E Envs were recognized efficiently by the pNAbs.

### Effect of crosslinking on the antigenicity of Env in SMALPs

The above study suggests that the 2-4 RM6 E Env in SMALPs samples conformations different from those associated with the membrane Env. Crosslinking has been shown to reduce HIV-1 Env conformational flexibility, decreasing the exposure of pNAb epitopes (33,37,71,88,96). We hypothesized that crosslinking the HIV-1 Env in the presence of BMS-806 prior to extraction from the membrane would help to preserve a pretriggered (State-1) conformation after solubilization in SMA. As the 2-4 RM6 E Env is lysine-rich (83), we chose the lysine-specific crosslinker, DTSSP. DTSSP crosslinks are able to be reduced, allowing crosslinked gp120 and gp41 subunits to be distinguished from uncleaved gp160 on Western blots. Membranes prepared from A549 cells expressing the 2-4 RM6 E Env were crosslinked with DTSSP in the presence of BMS-806. BMS-806-treated control membranes were mock treated without crosslinker in parallel. The membranes were solubilized with SMA and the 2-4 RM6 E Env was purified on Ni-NTA beads. After elution of Env, the binding of antibodies was studied by immunoprecipitation, as described above.

DTSSP crosslinking of the 2-4 RM6 E Env reduced recognition of the cleaved Env by the 19b anti-V3 pNAb, and decreased binding of the 19b and F240 pNAbs to the uncleaved gp160 Env (Fig. 9). The binding of bNAbs to the cleaved 2-4 RM6 E Env was not significantly affected by DTSSP treatment. The relatively inefficient recognition of the 2-4 RM6 E Env in SMALPs by the PGT145 and VRC03 bNAbs was not improved by DTSSP crosslinking prior to solubilization. These observations suggest that crosslinking may be useful in reducing the exposure of some pNAb epitopes that accompanies Env solubilization in SMA. However, crosslinking did not prevent the Env conformational changes associated with extraction into SMALPs that diminish recognition by the State-1-preferring PGT145 and VRC03 bNAbs.

**FIG 9.**
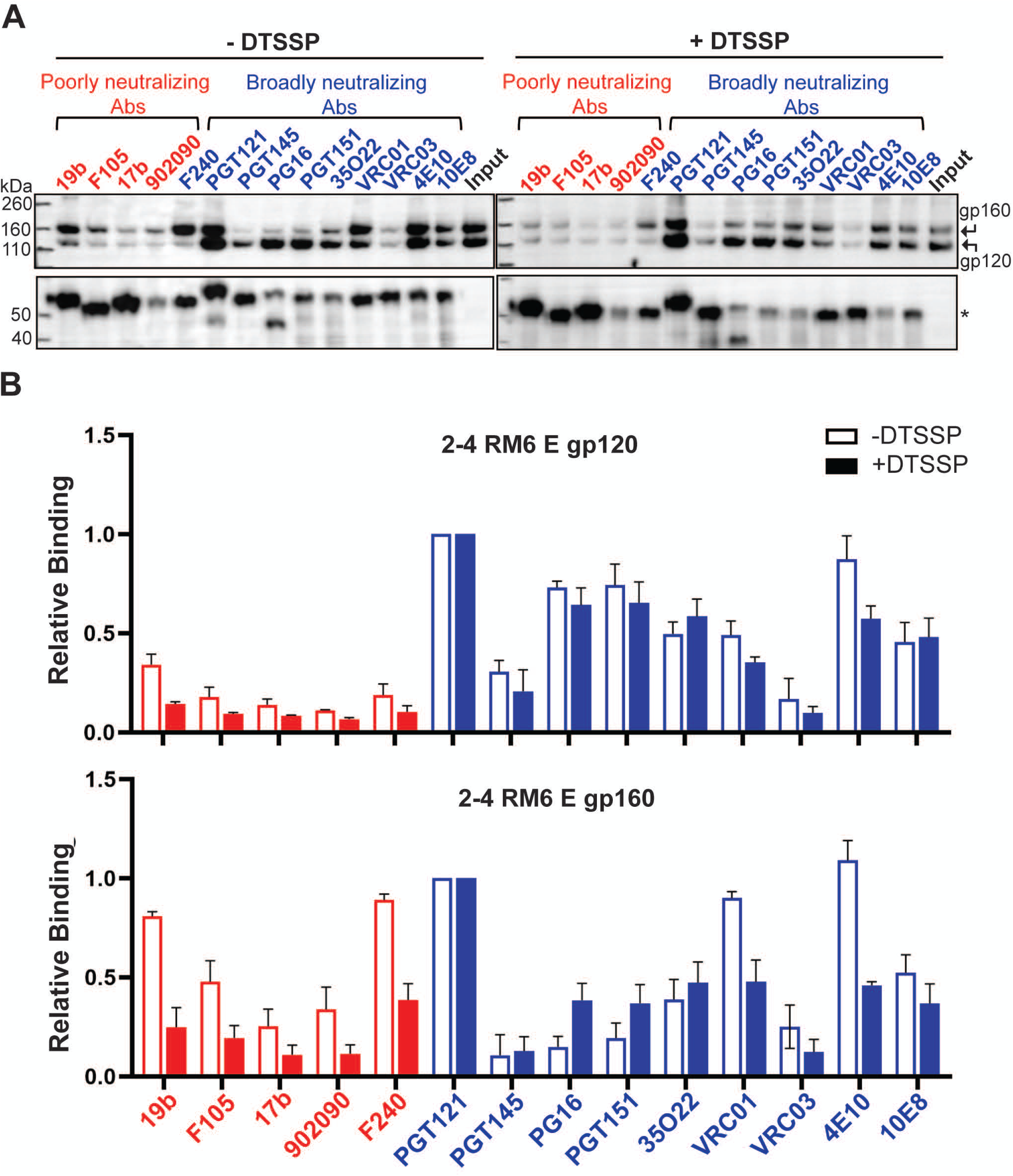
Effect of crosslinking on the antigenicity of 2-4 RM6 E Env-SMALPs prepared from purified membranes. (A) A549 cells were induced with doxycycline to express the 2-4 RM6 E Env. Cell membranes were purified and divided into two parts, one treated with 0.35 mM DTSSP crosslinker at room temperature for 45 minutes and the other mock-treated. DTSSP crosslinks are able to be reduced, allowing crosslinked gp120 and gp41 subunits to be distinguished from uncleaved gp160 on Western blots. The membranes were lysed in 1% SMA in the presence of 10 µM BMS-806 for 30 minutes at room temperature. The 2-4 RM6 E Env was purified by Ni-NTA affinity chromatography, aliquoted and incubated with the indicated antibodies together with Protein A-Sepharose beads for 1 h at 4°C. The precipitated proteins were analyzed on reducing SDS-polyacrylamide gels and Western blotted with a goat anti-gp120 antiserum (upper panels) and the human 4E10 anti-gp41 antibody (lower panels). The asterisk indicates the position of the heavy chain of the precipitating antibodies. (B) Quantification of gp120 and gp160 band intensity in A was performed in Fiji Image J (NIH). The intensities of the gp120 and gp160 bands were normalized to those of the respective bands precipitated by the PGT121 antibody. The means and standard deviations derived from two independent experiments are shown.

### Effect of pNAb counterselection on 2-4 RM6 E Env antigenicity

As the uncleaved Env is conformationally flexible and not functional (8,33–35,43,71), its removal from HIV-1 Env preparations would be desirable. The 19b V3-directed pNAb and the F240 pNAb against the gp41 ectodomain efficiently recognized the uncleaved Env (Fig. 10A). We tested whether the uncleaved gp160 Env could be preferentially removed from the 2-4 RM6 E Env preparation by counterselection with these pNAbs (97). Membranes from BMS-806-treated 2-4 RM6 E Env-expressing cells were crosslinked with DTSSP and used for the preparation of Env-SMALPs. After purification on Ni-NTA beads, the eluted Env-SMALPs were incubated with 19b and F240 pNAbs and Protein A-Sepharose beads. The antigenic profile of the 19b/F240-counterselected 2-4 RM6 E Env was evaluated as described above. Counterselection with the 19b and F240 pNAbs reduced the relative ratio of gp160:gp120 in the preparation (compare Input samples in Fig. 10A and B). The counterselected cleaved 2-4 RM6 E Env preparation was recognized by most of the bNAbs, except for the PGT145 bNAb against the V2 quaternary epitope at the trimer apex and the VRC03 CD4BS antibody (Fig. 10B). The recognition of the uncleaved gp160 by pNAbs was very inefficient following 19b/F240 counterselection. These results indicate that pNAb counterselection of BMS-806/DTSSP-treated Env-SMALPs can enrich the cleaved Env, which remains accessible to most bNAbs and less accessible to pNAbs.

**FIG 10.**
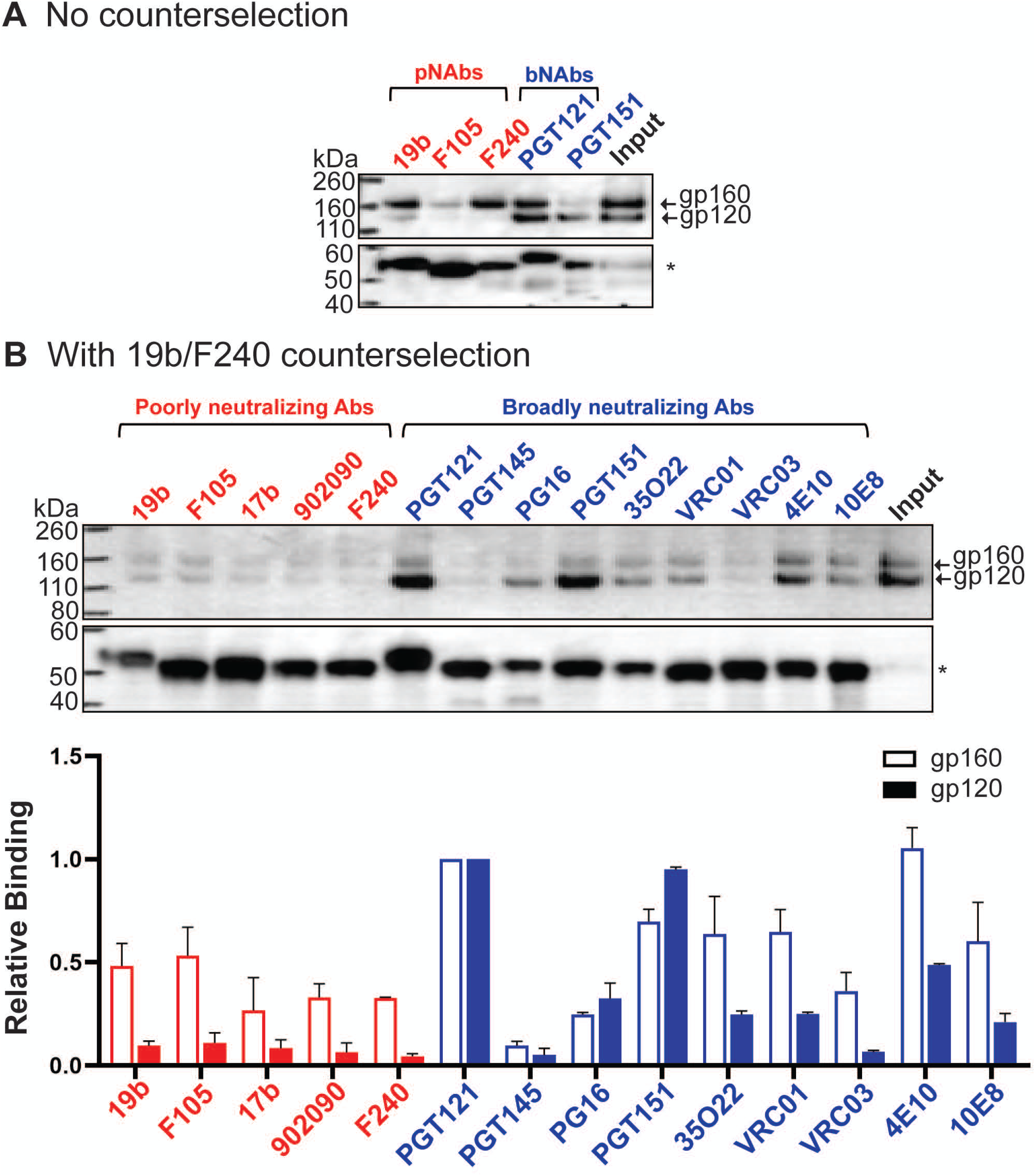
Antigenicity of Env purified using multiple measures to preserve Env conformation. (A) Cell membranes were purified from A549 cells induced with doxycycline to express the 2-4 RM6 E Env. The purified membranes were crosslinked with 0.35 mM DTSSP in the presence of 10 µM BMS-806 at room temperature for 45 minutes. The membranes were lysed in 1% SMA in the presence of 10 µM BMS-806 for 30 minutes at room temperature. The 2-4 RM6 E Env was purified from the membrane lysates by Ni-NTA affinity chromatography. In A, a fraction of the purified Env was saved as Input and the rest was used to evaluate the binding of the indicated antibodies, as described in the Figure 8 legend. In B, the purified Env was incubated with 40 µg/ml each of the 19b and F240 pNAbs together with Protein A-Sepharose beads for 30 minutes at room temperature. After 19b/F240 counterselection, the flow-through sample was aliquoted, with one fraction serving as the Input sample and the rest used to evaluate antigenicity, as described in the Figure 8 legend. The bar graph shows the relative binding to gp120 and gp160 normalized to the respective bands for the PGT121 antibody. The means and standard deviations from two experiments are reported.

## DISCUSSION

The metastability of the pretriggered (State-1) Env conformation, which is of great interest as a target for small-molecule virus entry inhibitors and vaccine-elicited antibodies, creates challenges for its purification and characterization. The cleaved Env in cell membranes exhibits an antigenic profile that strongly correlates with HIV-1 sensitivity to neutralization (43, 71). This observation suggests that the cleaved cell membrane Env closely approximates the functional pretriggered (State-1) Env conformation on the virion. In this study, we take several approaches to address the challenges for HIV-1 Env purification created by the requirement that Env must be extracted from its native membrane environment. Several studies characterizing phenotypes of HIV-1 Envs with alterations in the membrane-proximal external region (MPER) of gp41 have suggested a dependence of the State-1 conformation on Env association with the membrane (60–65). We evaluated the consequences of extracting HIV-1 Env from membranes on the level of cleavage, trimer integrity and antigenicity. In solubilizing Env, we took measures that, in the case of other integral membrane proteins, minimize conformational disruption (74-81,89,90,92,93). In addition to using a lysine-rich, State-1-stabilized Env variant treated with BMS-806 and a crosslinker, we explored the use of amphipathic copolymers that bypass the need for detergents, solubilizing membrane proteins directly in nanodiscs.

SMA efficiently extracted HIV-1 Env from cell membranes. The stability of Env trimers in SMALPs, particularly at room temperature, was superior to that in Cymal-5 or SDS/deoxycholate. Even without the State-1-stabilizing entry inhibitor, BMS-806, HIV-1 Env-SMALPs maintained subunit association for over 16 hours at room temperature. The stability of these amphipathic copolymer-Env complexes may be useful in the establishment of cell-free models of Env function or in tests of Env immunogenicity.

Our antigenicity analyses revealed differences between the 2-4 RM6 E Env in cell membranes and the Envs solubilized in SMA or Cymal-5. In the absence of BMS-806, V3, CD4BS and CD4i pNAb epitopes are exposed in the Env-SMALPs and in Cymal-5-solubilized Env, indicating sampling of non-State-1 conformations. Without BMS-806, recognition of the quaternary V2 epitope at the trimer apex by PGT145 and the VRC03 CD4BS epitope was relatively inefficient for the Cymal-5-solubilized Env and Env-SMALPs. As the PGT145 and VRC03 bNAbs demonstrate some preference for the functional State-1 conformation (68, 69), weak recognition of these solubilized Envs is consistent with disruption of the State-1 conformation. The conformational effects associated with extraction of the 2-4 RM6 E Env from membranes could be partially but not completely mitigated by pre-treatment of the membrane Env with BMS-806 and a crosslinker. Treatment of the membrane 2-4 RM6 E Env with BMS-806 and DTSSP decreased the exposure of pNAb epitopes in the Env-SMALPs and enhanced recognition by some, but not all, bNAbs. In particular, PGT145 and VRC03 binding was still weak, suggesting that the 2-4 RM6 E Env in the BMS-806/DTSSP-treated Env-SMALPs assumes a conformation that resembles, but is still distinct from, State 1.

What conformations are assumed by the solubilized Envs in Cymal-5 micelles or SMALPs? Subtle differences between these solubilized Env complexes were detected in the gp41 MPER. The 4E10 and 10E8 bNAbs against the gp41 MPER recognized the Cymal-5-solubilized 2-4 RM6 E Env more efficiently than the SMA-solubilized Env, in the absence of BMS-806. Differential recognition by these antibodies presumably reflects structural differences in the MPER associated with these distinct methods of Env solubilization. Other than these differences in recognition by MPER-directed bNAbs, the antigenicity of the Cymal-5-solubilized and SMA-solubilized 2-4 RM6 E Envs was similar. The correlation between the antigenic profiles of the 2-4 RM6 E Env solubilized in Cymal-5 and SMA indicates that membrane extraction without BMS-806 results in similar non-State-1 Env conformations in these two preparations (Fig. 11). Of interest, State 2 has been suggested to represent a default intermediate conformation assumed when the State-1 conformation of the membrane Env is disrupted (68, 69). Env solubilization may likewise consign Env to a default conformation resembling State 2, a model consistent with the smFRET profile of a Cymal-5-solubilized uncleaved Env (71). Very recent cryo-EM structures of cleaved HIV-1 Envs in SMALPs reveal asymmetric trimers (98); the distinct pattern of opening angles among the protomers resembles that seen for the uncleaved Env solubilized in Cymal-5 and amphipol A8-35 (71). These structures represent reasonable candidates for the default State-2 Env conformation that, at least upon binding CD4, is thought to be asymmetric (99). The Env-SMALP structures suggest that State 1-to-State 2 transitions may involve breaking trimer symmetry coupled with trimer tilting in the membrane (98). Together, these observations support a model in which disruption of the metastable symmetric State-1 conformation by Env solubilization and loss of membrane interactions leads to an asymmetric trimer in the default intermediate (State-2) conformation. Changes in Env structure that are coupled to the breaking of symmetry result in the stabilization of State-2 (98), explaining the difficulty of reverting State 2 to State 1 in the absence of a membrane (59, 85).

**FIG 11.**
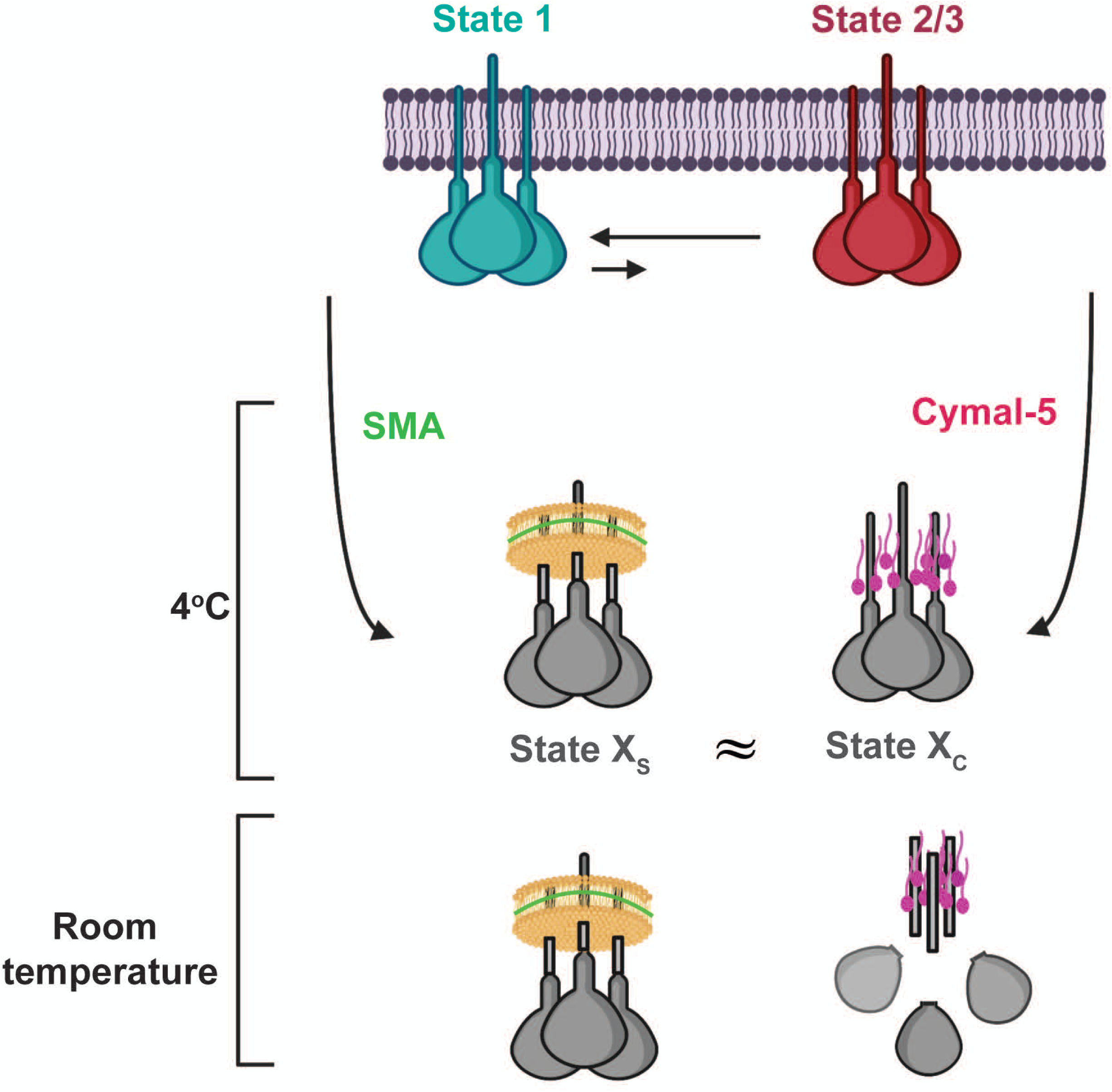
A model for the effect of extraction from membranes on HIV-1 Env conformation. The cleaved membrane Env of primary HIV-1 mainly occupies a pretriggered (State-1) conformation. Upon solubilization of the membrane proteins by SMA or Cymal-5 at 4°C, the extracted Envs assume conformations (States X_S_ and X_C_, respectively) that differ from State 1. The conformations of the Envs in SMALPs and Cymal-5 resemble one another, with differences mainly confined to the membrane-proximal Env elements. The resemblance of the Envs solubilized by distinct approaches supports the existence of a default Env conformation that is assumed when State 1 is destabilized by the loss of the membrane. BMS-806 can bring the solubilized Envs closer to a State-1 conformation, but BMS-806-treated, solubilized Envs still differ from State 1. When the temperature is raised to room temperature, Envs in SMALPs are more stable than Envs in detergents like Cymal-5.

A more complete understanding of the differences between State 1 and State 2 could assist the design of small-molecule entry inhibitors that prematurely activate or block Env transitions between these conformations (84-87,100). The antigenic differences that we identified between the solubilized cleaved Envs (potentially in State-2-like conformations) and the cleaved membrane Env (mostly in a State-1 conformation) may affect the efficacy of vaccine immunogens. A recent antigenicity analysis has suggested that only modest differences exist between the State-1 and State-2 Env conformations, although failure to distinguish cleaved and uncleaved Envs in that study may have diminished the ability of some of the antibodies to discriminate between these conformations (100). The results reported herein will guide future efforts to stabilize the State-1 Env further and to improve methods to solubilize HIV-1 Env with minimal conformational disruption.

## MATERIALS AND METHODS

### HIV-1 Env constructs

The wild-type (wt) HIV-1_AD8_, 2-4 RM6 E and AE.2 Envs were coexpressed with the Rev protein, using the natural arrangement of HIV-1 *env* and *rev* sequences (60, 83). The Asp 718 (Kpn I) – Bam HI *env* fragments encoding the above Envs were inserted into the corresponding sites of the pSVIIIenv plasmid expressing the HIV-1_HXBc2_ Env and Rev. Thus, these Envs contain a signal peptide and part of the cytoplasmic tail from the HIV-1_HXBc2_ Env. A GGGHHHHHH (His_6_) tag was added to the carboxyl terminus of all three Envs. Compared with the HIV-1_AD8_ Env, the 2-4 RM6 E Env has two State-1-stabilizing changes, Q114E and Q567K, and several additional lysine substitutions to improve crosslinking efficiency (R166K, R178K, R315K, R419K, R557K, R633K, Q658K, A667K and N677K) (83). Compared with the 2-4 RM6 E Env, the AE.2 Env has one additional State-1-stabilizing change, A582T, and an additional lysine substitution, R252K (83).

### SMA and DIBMA

SMA (2:1) was purchased from Cray Valley and hydrolyzed as described (77). SMA and DIBMA variants were purchased from Cube Biotech.

### Reagents

BMS-378806 (herein called BMS-806) was purchased from Selleckchem. Cymal-5 and Cymal-6 were purchased from Anatrace, RIPA buffer from ThermoFisher and Superflow Ni-NTA from Bio-Rad.

### Expression of HIV-1 Envs

Human A549 cells inducibly expressing the wt HIV-1_AD8_ Env, the 2-4 RM6 E Env and the AE.2 Env were established as described (43). A549 cells constitutively expressing the reverse tet transactivator (rtTA) were transduced with HIV-1-based lentivirus vectors expressing Rev and the above Envs. The vector transcribes a bicistronic mRNA comprising *rev* and *env* and two selectable marker genes (puromycin-T2A-enhanced green fluorescent protein (EGFP)) fused in-frame with a T2A peptide-coding sequence. In the transduced cells, Env expression is controlled by the Tet-Responsive Element (TRE) promoter and tet-on transcriptional regulatory elements. Env expression is induced by treating the cells with 2 ug/ml doxycycline. The Env-expressing cells were enriched by fluorescence-activated cell sorting for the co-expressed EGFP marker. These polyclonal A549 cell lines were used as sources of Env for the studies reported herein.

### Cell lines

The A549 cells inducibly expressing the wt HIV-1_AD8_, 2-4 RM6 E and AE.2 Envs were grown in DMEM/F12 supplemented with 10% FBS, L-glutamine and penicillin-streptomycin. All cell culture reagents are from Life Technologies.

### Antibodies

Antibodies against HIV-1 Env were kindly supplied by Dr. Dennis Burton (Scripps), Drs. Peter Kwong and John Mascola (Vaccine Research Center, NIH), Dr. Barton Haynes (Duke), Dr. Hermann Katinger (Polymun), Dr. James Robinson (Tulane) and Dr. Marshall Posner (Mount Sinai Medical Center). In some cases, anti-Env antibodies were obtained through the NIH AIDS Reagent Program. Antibodies for Western blotting include goat anti-gp120 polyclonal antibody (ThermoFisher) and the 4E10 human anti-gp41 antibody (Polymun). An HRP-conjugated goat anti-human IgG (Santa Cruz) and an HRP-conjugated goat anti-rabbit antibody (Santa Cruz) were used as secondary antibodies for Western blotting.

### Immunoprecipitation of cell-surface HIV-1 Env

Doxycycline-induced A549-Env cells were washed twice with washing buffer (1x PBS + 5% FBS). The cells were then incubated with 5 µg/ml antibody for one hour at 4°C. After washing four times in washing buffer, the cells were lysed in NP-40 lysis buffer (1% NP-40, 0.5 M NaCl, 10 mM Tris, pH 7.5) for five minutes on ice. The lysates were cleared by centrifugation at 13,200 x g for 10 minutes at 4°C, and the clarified supernatants were incubated with Protein A-Sepharose beads for one hour at room temperature. The beads were pelleted (1,000 rpm x 1 min) and washed three times with wash buffer (20 mM Tris-HCl (pH 8.0), 100 mM (NH_4_)_2_SO_4_, 1 M NaCl and 0.5% NP-40). The beads were suspended in 2x lithium dodecyl sulfate (LDS) sample buffer, boiled and analyzed by Western blotting using 1:2000 goat anti-gp120 polyclonal antibody (ThermoFisher) and 1:2000 HRP-conjugated rabbit anti-goat IgG (ThermoFisher). The HIV-1 gp41 Env was analyzed by Western blotting with the 4E10 anti-gp41 antibody and HRP-conjugated goat anti-human IgG (Santa Cruz).

For analysis of total Env expression in the cells, clarified cell lysates were prepared from cells that were induced with doxycycline but not incubated with antibodies. The clarified cell lysates were analyzed by Western blotting as described above and serve as the Input samples.

### Membrane purification and DTSSP crosslinking

A549 cells expressing the wt HIV-1_AD8_ or 2-4 RM6 E Envs were incubated with 5 mM EDTA in 1x PBS at 37°C until the cells detached from the tissue-culture plates. The cells were pelleted and resuspended in 1x PBS. Cells were spun down at 1500 x g for 10 minutes. The supernatants were removed and homogenization buffer (10 mM Tris HCl (pH 7.5), 250 mM sucrose, 1 mM EDTA, 1x protease inhibitor cocktail) was added to the cell pellet. The cells were transferred into a glass Dounce homogenizer and homogenized with 250 strokes on ice. The homogenate was spun at 1000 x g for 10 minutes at 4°C. The supernatants were centrifuged at 10,000 x g for 10 minutes at 4°C. The supernatants were spun again at 100,000 x g for 35 minutes at 4°C. The pellet represented the purified membrane fraction.

For DTSSP crosslinking, membranes were washed in 1x PBS and centrifuged at 100,000 g for 25 minutes. Membranes were resuspended in 1x PBS and crosslinked with 0.35 mM DTSSP at 4°C for 45 minutes.

### Immunoprecipitation of cell-membrane HIV-1 Envs

Purified cell membranes from Env-expressing A549 cells were incubated with 5 ug/ml antibody for one hour at 4°C, in either the absence or presence of 10 µM BMS-806. The membranes were pelleted at 13,200 x g for 10 minutes at 4°C. The pelleted membranes were lysed in 1% NP-40 with or without BMS-806 for 5 minutes on ice. The membrane lysates were then incubated with Protein A-Sepharose beads for one hour at 4°C. The precipitated proteins were subjected to Western blotting with a goat anti-gp120 antiserum or the human 4E10 anti-gp41 antibody, as described above.

For the Input sample, an aliquot of the cell membrane preparation was lysed in 1% NP-40 and subjected to Western blotting as described above.

### Solubilization of Env from membranes

For extraction of Env from membranes, the purified membrane fractions were resuspended in the following buffers:

SMA solubilization buffer (20 mM Tris-HCl, pH 8.0, 100 mM (NH_4_)_2_SO_4_, 250 mM NaCl 1% SMA, 1x protease inhibitor cocktail (Roche));

DIBMA solubilization buffer (20 mM Tris-HCl, pH 8.0, 100 mM (NH_4_)_2_SO_4_, 250 mM NaCl, 1% DIBMA, 1x protease inhibitor cocktail (Roche));

Cymal-5 solubilization buffer (20 mM Tris-HCl, pH 8.0, 100 mM (NH_4_)_2_SO_4_, 250 mM NaCl, 1% Cymal-5, 1x protease inhibitor cocktail (Roche)); and

RIPA buffer (ThermoFisher) (25 mM Tris-HCl, pH 7.6, 150 mM NaCl, 1% NP-40,1% sodium deoxycholate, 0.1% SDS, 1x protease inhibitor cocktail (Roche)).

Solubilized Envs were analyzed on reducing and non-reducing gels. For comparison of SMA and DIBMA variants, gels were silver stained using the Pierce Silver stain kit (Thermo Scientific) or Western blotted as described above.

### Solubilization of Env from cells

Env-expressing cells were induced with 2 ug/ml of doxycycline for two days to express HIV-1 Envs and detached by incubating with 5 mM EDTA at 37°C. Cells were aliquoted and lysed with the solubilization buffers described above at either 4°C or room temperature.

### Association of gp120 with solubilized Env complexes

The noncovalent association of gp120 with solubilized Env complexes was studied by precipitating the proteins using the carboxy-terminal His_6_ tag. Clarified cell lysates were prepared from Env-expressing A549 cells as described above. Aliquots of the lysates were saved for Western blotting to detect the gp160, gp120 and gp41 glycoproteins in the Input sample. The bulk of the lysates was incubated for 1 h or 16 h at 4°C or room temperature, in some cases in the presence of 10 µM BMS-806. The lysates were then incubated for 1 h with nickel-nitriloacetic acid (Ni-NTA) beads (Qiagen) at the indicated temperatures (4°C or room temperature), in some cases with 10 µM BMS-806. The beads were pelleted (1,000 rpm for 1 min at 4°C or room temperature), washed three times at 4°C or room temperature with wash buffer (20 mM Tris-HCl (pH 8.0), 100 mM (NH_4_)_2_SO_4_, 1 M NaCl, 30 mM imidazole), boiled in 2x LDS sample buffer, and analyzed by Western blotting as described above. The association of gp120 with the Env complex for each solubilization buffer was calculated as follows: (gp120/gp160)_precipitated_ + (gp120/gp160)_Input_.

### Immunoprecipitation of solubilized Envs from cell lysates

A549 cells were induced with doxycycline to express the 2-4 RM6 E Env. Forty-eight hours later, the cells were lysed with either Cymal-5 or SMA solubilization buffer, as described above. The 2-4 RM6 E Env was incubated with Ni-NTA beads at 4°C for 1.5 hours. After incubation, the mixture was applied to an Eco-column (Biorad). The beads were washed with 30 bed volumes of washing buffer (20 mM Tris-HCl (pH 8.0), 100 mM (NH_4_)_2_SO_4_, 1 M NaCl, 30 mM imidazole), and eluted with 10 bed volumes of elution buffer (20 mM Tris-HCl (pH 8.0), 100 mM (NH_4_)_2_SO_4_, 250 mM NaCl, 250 mM imidazole). The purified Env was aliquoted and incubated with 10 µg/ml antibodies together with Protein A-Sepharose beads for 1 h at 4°C. For the samples with BMS-806, 10 µM BMS-806 was added before cell lysis and remained in all the following steps. An aliquot without added antibody/beads served as the Input sample. The precipitated proteins and Input sample were Western blotted as described above.

### Antigenicity of Envs solubilized from purified membranes

Cell membranes were prepared from A549 cells expressing the 2-4 RM6 E Env as described above. In some cases, the resuspended cell membranes were treated with 0.35 mM DTSSP crosslinker at room temperature for 45 minutes. In some experiments, BMS-806 at 10 µM concentration was added to the membranes at the time of crosslinking. The membranes were then lysed in SMA solubilization buffer containing 10 µM BMS-806 for 30 minutes at room temperature. The 2-4 RM6 E Env was incubated with Ni-NTA beads at 4°C for 1.5 hours. After incubation, the mixture was applied to an Eco-column (Biorad). The Ni-NTA beads were washed with 30 bed volumes of washing buffer (20 mM Tris-HCl (pH 8.0), 100 mM (NH_4_)_2_SO_4_, 1 M NaCl, 30 mM imidazole), and eluted with 10 bed volumes of elution buffer (20 mM Tris-HCl (pH 8.0), 100 mM (NH_4_)_2_SO_4_, 250 mM NaCl, 250 mM imidazole). For the experiments where counterselection with pNAbs was employed, the purified 2-4 RM6 E Env was incubated with 40 µg/ml of 19b and 40 µg/ml of F240 antibodies and 100 µl of Protein A-Sepharose at room temperature for 30 minutes. The mixture was applied to an Eco-column (Biorad) and the purified 2-4 RM6 E Env was collected in the flow-through.

## ACKNOWLEDGMENTS

We thank Ms. Elizabeth Carpelan for manuscript preparation. Antibodies against HIV-1 were kindly supplied by John C. Kappes (University of Alabama at Birmingham), Dennis Burton (Scripps), Peter Kwong and John Mascola (Vaccine Research Center NIH), Barton Haynes (Duke University), Hermann Katinger (Polymun), James Robinson (Tulane University), and Marshall Posner (Mount Sinai Medical Center). We thank the NIH HIV Reagent Program for providing additional reagents.

This work was supported by grants from the National Institutes of Health (grant nos. AI 145547, AI 124982, AI 150471, AI 129017 and AI 164562), a grant from Gilead Sciences, and by a gift from the late William F. McCarty-Cooper.

We declare no conflicts of interest.

